# Left Atrial Strain as a Predictor of Cardiac Dysfunction in a Murine Model of Pressure Overload

**DOI:** 10.1101/2024.09.05.611376

**Authors:** John P. Salvas, Thomas Moore-Morris, Craig J. Goergen, Pierre Sicard

## Abstract

**Aim:** Left atrial (LA) strain is emerging as a valuable metric for evaluating cardiac function, particularly under pathological conditions such as pressure overload. This preclinical study investigates the predictive utility of LA strain on cardiac function in a murine model subjected to pressure overload, mimicking pathologies such as hypertension and aortic stenosis.

**Methods:** High resolution ultrasound was performed in a cohort of mice (n=16) to evaluate left atrial and left ventricular function at baseline and 2- and 4-weeks after transverse aortic constriction (TAC). Acute adaptations in cardiac function were assessed in a subgroup of mice (n=10) with 3-days post TAC imaging.

**Results:** We report an increase in LA max volume from 11.0 ± 4.3µL at baseline to 26.7 ± 16.7µL at 4 weeks (*p*=0.002) and a decrease in LA strain from 19.6 ± 4.8% at baseline to 10.1 ± 6.3% at 4 weeks (*p*=0.006). In the acute phase, LA strain dysfunction was present at 3-days (*p*<0.001) prior to alterations in LA volume (*p*=0.856) or left ventricular (LV) ejection fraction (*p*=0.120). LA strain correlated with key indicators of cardiac performance including left ventricular (LV) ejection fraction (r=0.563, *p*<0.001), longitudinal strain (r=-0.643, *p*<0.001) and strain rate (r=0.387, *p*=0.007). Furthermore, markers of atrial structure and function including LA max volume (AUC=0.858, *p*<0.001), ejection fraction (AUC=0.901 *p*<0.001), and strain (AUC=0.878, *p*<0.001) all predicted LV dysfunction.

**Conclusion:** LA strain and function assessments provide a reliable, non-invasive method for early detection and prediction of cardiac dysfunction in a model of pressure overload.

## INTRODUCTION

Left atrial (LA) remodeling are common in various cardiovascular conditions, including atrial fibrillation, diastolic dysfunction, and heart failure (1,2). Monitoring LA evolution during cardiovascular disease has emerged as a valuable biomarker for risk stratification and ongoing risk assessment (3,4). Beyond just measuring the size, assessing LA function dynamics provides critical insights into the heart’s structural and functional adaptations, particularly in the characterization of left ventricular (LV) diastolic function (5). This is especially important in heart failure where LA dysfunction can significantly affect overall cardiac output (6). In particular, pressure overloaded conditions, such as those induced by uncontrolled hypertension or aortic stenosis, place a significant burden on the LV, which in turn impacts the LA. As the LV faces increased afterload, the LA must exert greater effort to maintain efficient blood flow and support ventricular filling. This increased workload on the LA can lead to chronic alterations in its structure and function (7,8). Recently, LA strain, a measure of the deformation of the LA myocardium during the cardiac cycle, has gained attention for its potential to provide deeper insights into atrial function (9). In clinical settings, early changes in LA mechanics are often associated with the severity of heart failure and other adverse cardiovascular outcomes (10,11). However, despite its clinical relevance, the measurement of LA strain in murine experimental models of heart failure remains underexplored (12,13). Here, we used advanced echocardiographic techniques to precisely quantify LA strain and examined its relationship with other indices of LV dysfunction following acute and chronic transverse aortic constriction (TAC). Overall, our findings suggest that acute altered LA strain is predictive of cardiac dysfunction in pressure overload.

## RESULTS

### Ultrasound Reveals Progressive Left Atrial Remodeling Following TAC

High-resolution ultrasound was used to assess left atrial anatomical and functional activity two and four weeks after TAC (Figure 1). We performed 2D scans in the short-axis view at the base of the heart to obtain a clear view of the left atrium (Figure 1B-C). Consistent with previous reports (14), the LA showed signs early structural remodeling due to cardiac pressure overload, indicated by a progressive increase in LA maximum volume (baseline: 11.0 ± 4.3µL, 2W TAC: 18.0 ± 11.4µL; 4w TAC: 26.7 ± 16.7µL, *p*=0.002; Figure 1D). In addition, four-dimensional (4D) ultrasound assessments of LA max volume showed a similar significant increase in max volume from baseline to 4 weeks post TAC (*p<*0.001) and correlated with 2D volume assessments (r=0.775, *p<*0.0001; Supplemental Figure 1). Atrial volume enlargement was associated with atrial interstitial fibrosis, evidenced by an increase in Collagen type I^+^ area (baseline: 4.8 ± 1.1%, 4w TAC 10.5 ± 1.7%; Figure 1E-G).

**Figure 1:**
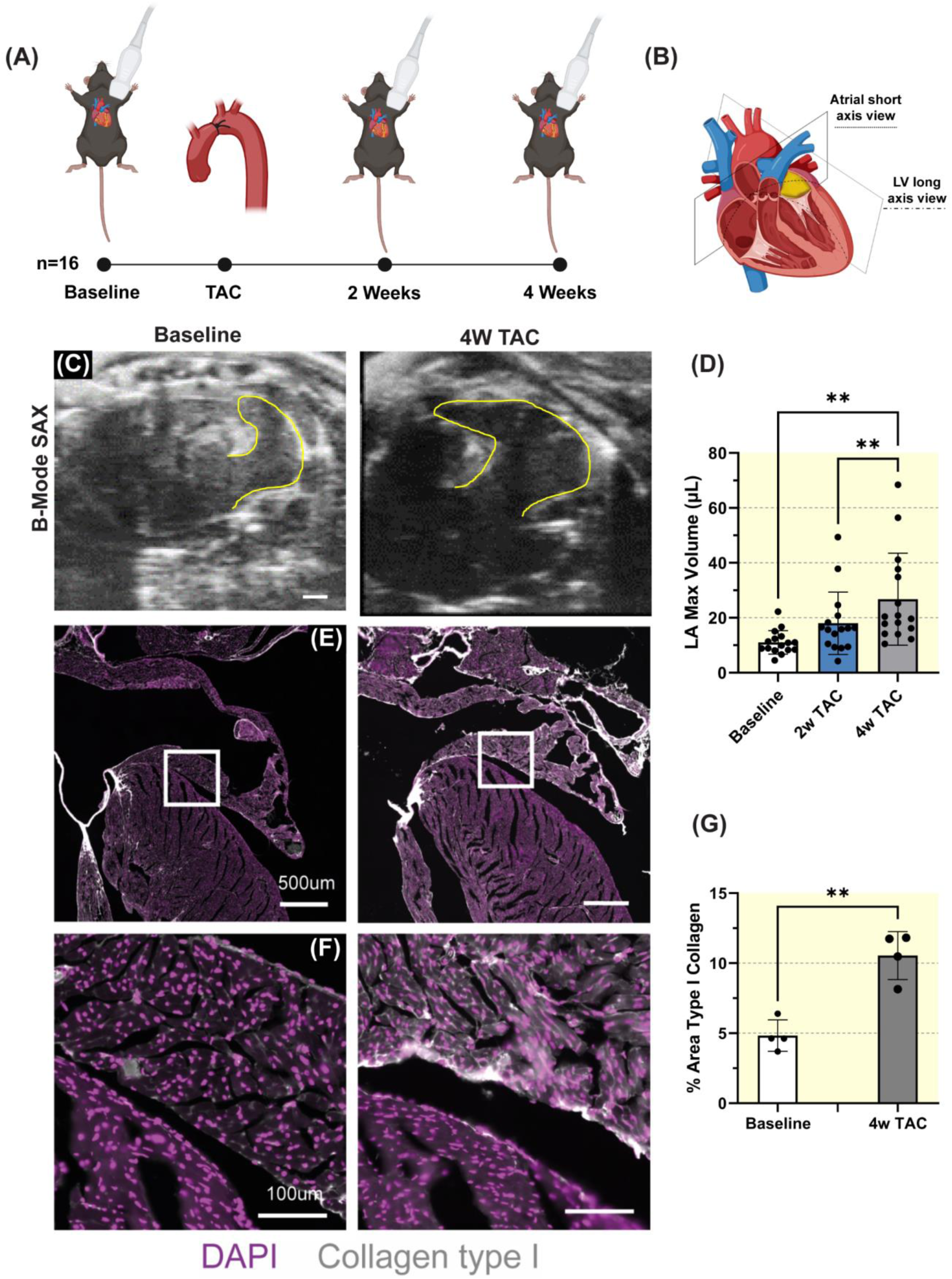
Experimental design and left atrial histology. **(A)** Study design and timeline. **(B)** Representation of left atrial short-axis and left ventricular long-axis imaging. **(C)** Representative short-axis ultrasound images of the left atrium at baseline and 4-weeks post-TAC procedure. Scale bar represents 1mm. **(D)** Left atrial maximum volume. Sample size n=16. Results expressed as mean ± SD. Statistical comparison conducted with one-way ANOVA with Tukey’s post hoc multiple comparisons tests. **(E)** Representative histologic cross sections of the left atrium and atrial appendage at baseline and 4-weeks post-TAC. **(F)** Higher magnification histologic cross sections of the left atrium at baseline and 4-weeks post-TAC. Immunofluorescence staining shows type I collagen (grayscale) and DAPI/nuclei (violet). **(G)** Left atrial fibrosis quantification in baseline and 4 weeks post-TAC animals. Sample size n=4. Results expressed as mean ± SD. Statistical comparison conducted with non-paired studentized t-test.

### Pressure Overload Leads to Sustained Decline in Left Atrial Strain assessed by speckle tracking

Next, we tested the hypothesis that pressure overload decreases LA function. Using a specific 2D basal short-axis view and a speckle tracking analysis allows for the calculation of LA function parameters including ejection fraction and strains (Figures 2A-B). LA ejection fraction (EF) significantly decreased from 50.5 ± 7.8% at baseline to 29.9 ± 12.9% at 2 weeks post-TAC (*p*<0.001) and remained consistently decreased at 4 weeks (31.4 ± 14.5, *p*=0.928; Figure 2C). Additionally, LA fractional area changes and circumferential strain showed similar trends with significant decreases from baseline to 2 weeks and non-significant changes from 2- to 4-weeks post TAC (Figure 2D-E). LA longitudinal strain was 19.6 ± 4.8% at baseline and significantly decreased to 8.2 ± 4.7% at 2 weeks (*p*<0.0001) and remained consistently decreased at 4 weeks compared to 2 weeks (10.1 ± 6.3%, *p*=0.589; Figure 2F).

**Figure 2:**
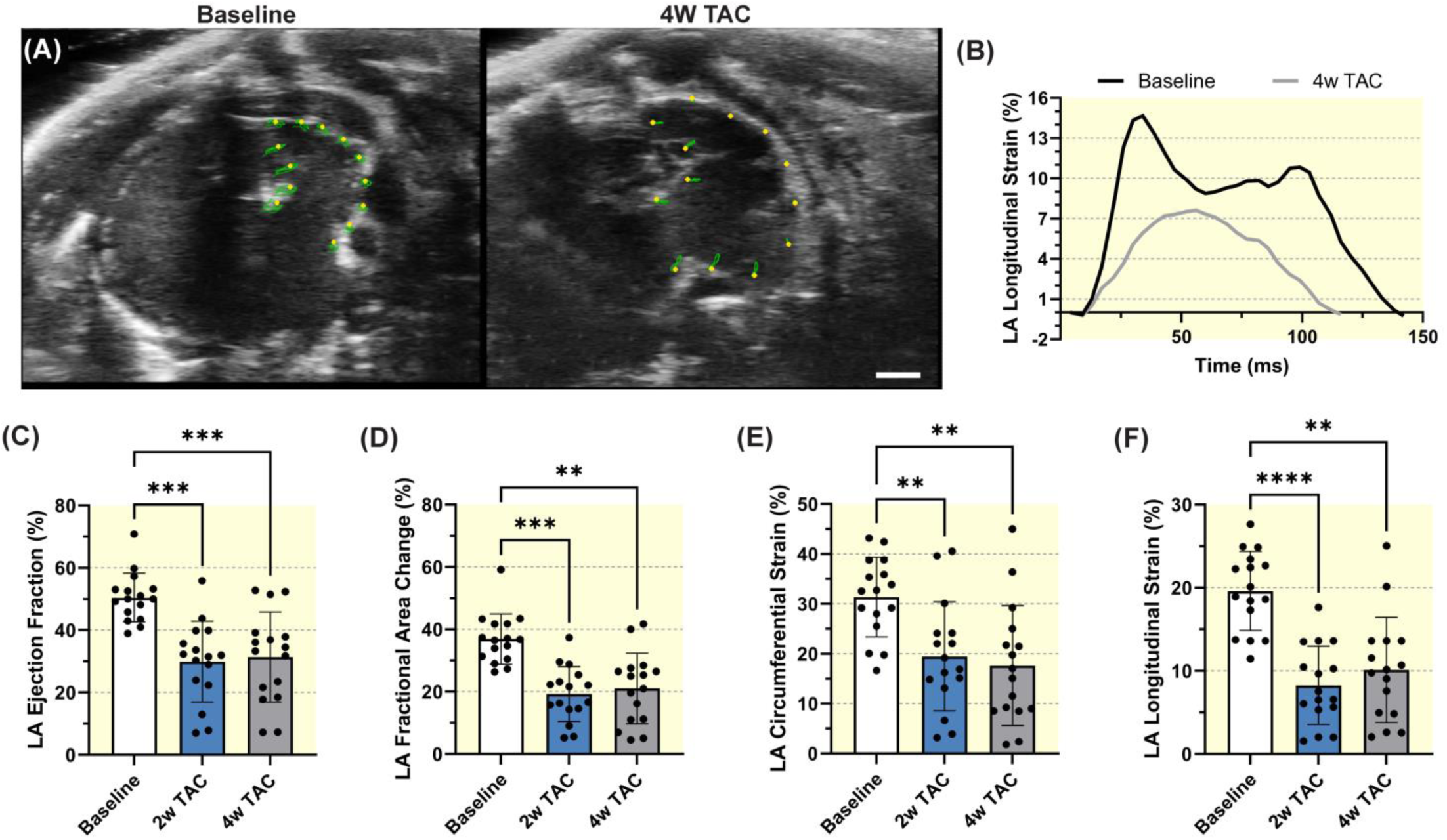
Speckle-tracking analysis of left atrial Function in a pressure overload model. **(A)** Representative baseline and 4 weeks post-TAC example short-axis views of the left atrium in the VevoStrain 2.0 software with yellow points representing the atrial myocardial wall tracking and green lines representing the myocardial motion over the cardiac cycle. Scale bar represents 1mm. **(B)** Representative global longitudinal strain curve for the left atrium at baseline and 4 weeks post-TAC. Left atrial **(C)** ejection fraction, **(D)** fractional area change, **(E)** circumferential strain, and **(F)** longitudinal strain. Sample size n=16 for each timepoint. Results expressed as mean ± SD. Statistical comparisons conducted with one-way ANOVA with Tukey’s post hoc multiple comparisons tests.

### Early Left Atrial Strain Dysfunction Predicts Late Left Ventricular Failure

We then evaluated how LA strain dysfunction associates with heart failure development. After TAC, animals displayed signs of cardiac decompensation indicated by significant reductions in LV contractile systolic and diastolic function (Figures 3A-D). Interestingly, LV EF was 61.5 ± 6.2% at baseline and progressively decreased from 52.4 ± 2.8% at 2 weeks to 42.2 ± 8.4% at 4 weeks (*p*<0.001) while LV longitudinal strain was –18.1 ± 2.1% at baseline and remained consistently decreased at -13.1 ± 2.6% and –11.7 ± 3.1% (*p*=0.238) at 2- and 4-weeks post TAC, respectively. LV early diastolic strain rate followed a similar trend with a significant reduction from baseline to 2 weeks (*p*=0.014) proceeded by no significant change at 4 weeks (*p*=0.287). Furthermore, LA strain measurements were significantly correlated with ventricular systolic function (ejection fraction: r=0.563, *p*<0.0001; strain: r=-0.643, *p*<0.0001) and diastolic function over time (early diastolic strain rate: r=0.387, *p*=0.007; Figure 3D-G). Further correlation between LA and LV function parameters seen in Supplemental Figure 2A. Thus, we confirm that LA strain correlates with LV dysfunction.

**Figure 3:**
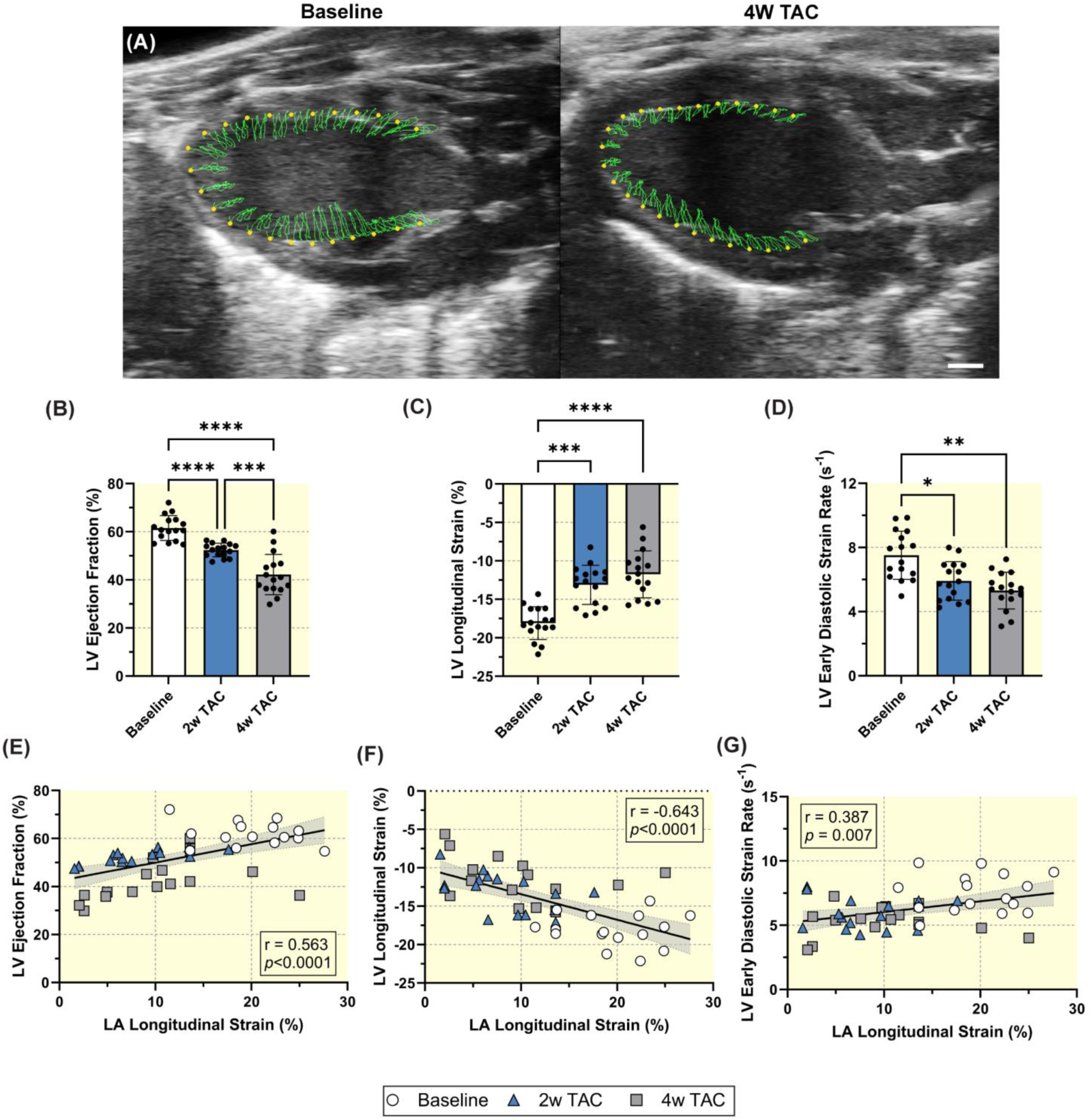
Left atrial function correlates with Left ventricular function. **(A)** Representative baseline and 4 weeks post-TAC parasternal long-axis views of the left ventricle in the VevoStrain 2.0 software with yellow points representing the ventricular myocardial wall tracking and green lines representing the myocardial motion over the cardiac cycle. Scale bar represents 1mm. Left ventricular **(B)** ejection fraction, **(C)** longitudinal strain, and **(D)** early diastolic strain rate. Sample size n=16 for each timepoint. Results expressed as mean ± SD. Statistical comparisons conducted with one-way ANOVA with Tukey’s post hoc multiple comparisons tests. Correlation of left ventricular **(E)** ejection fraction, **(F)** longitudinal strain and **(G)** early diastolic strain rate versus left atrial longitudinal strain. Sample size is n=48 which includes measurements from all timepoints. Pearson correlation coefficients and corresponding *p*-values displayed.

Moreover, we investigated the hypothesis that LA strain serves as a useful biomarker for identifying LV dysfunction via the correlation between LA function 2-weeks after TAC and LV function 4-weeks after TAC (Figure 4A-C). LA EF significantly correlated with both LV longitudinal strain (r=-0.546, *p=*0.032) and strain rate (r=0.504, *p*=0.047). Early LA strain also correlated with LV longitudinal strain (r=-0.553, *p*=0.026) at 4 weeks. Further correlations between early LA and late LV dysfunction are shown in Supplemental Figure 2B. Significant correlations were found, suggesting that early LA dysfunction quantified by LA strain predicted late LV dysfunction. Additionally, we further tested that LA dysfunction could serve as a useful imaging biomarker by discriminating between low and high LV EF groups, based on the median LV EF of 53.7%. The ROC curves illustrate the diagnostic performance of LA max volume (AUC=0.858), LA EF (AUC=0.901), and LA longitudinal strain (AUC=0.878) in differentiating between TAC mice with varying levels of LV EF. All parameters of LA function were able to significantly discriminate between low and high LVEF with *p*<0.0001 (Figure 4D).

**Figure 4:**
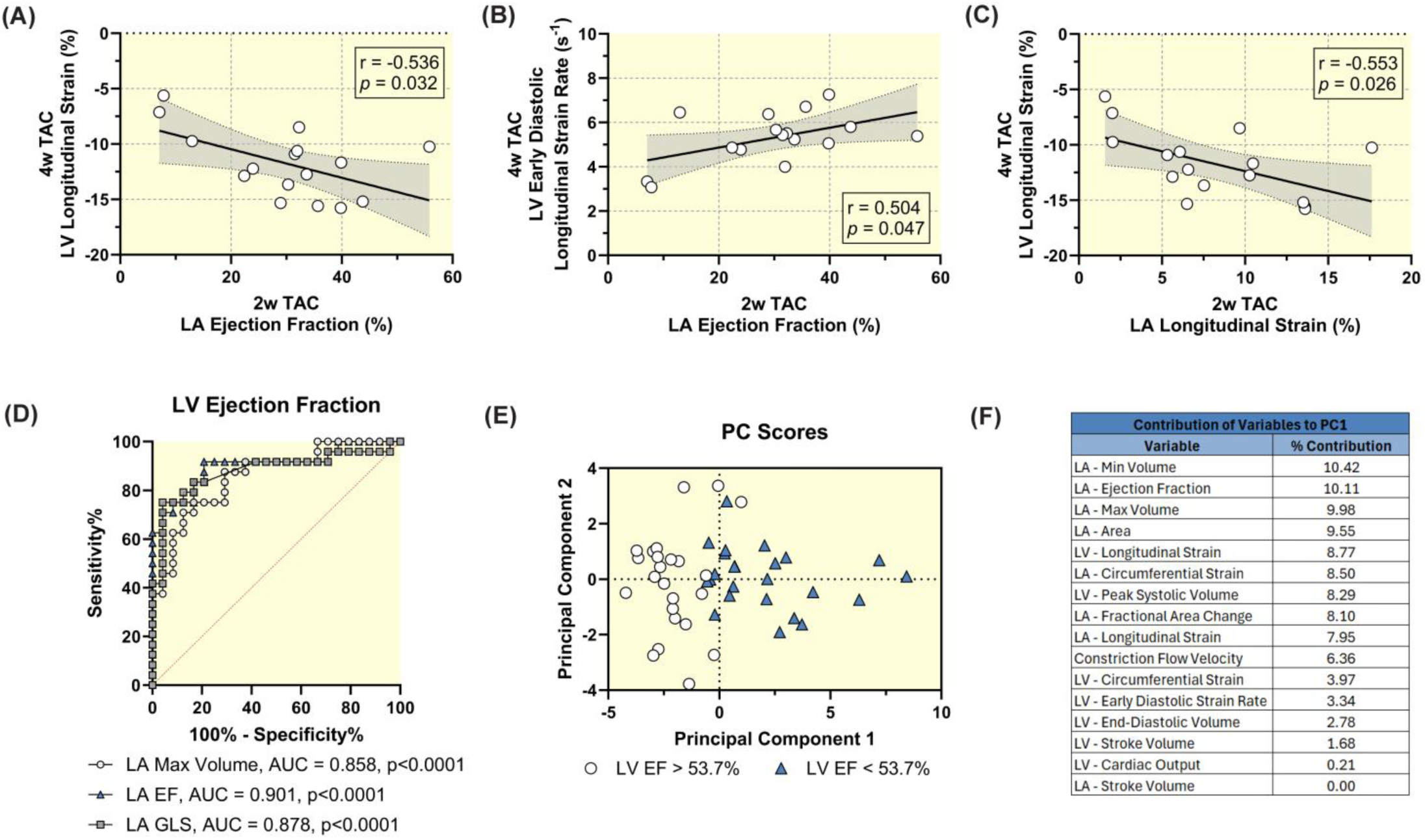
Early left atrial dysfunction predicts ventricular dysfunction. Pearson correlation of 4-week post-TAC left ventricular **(A)** longitudinal strain and **(B)** early diastolic strain rate verses 2-week post-TAC left atrial ejection fraction. **(C)** Pearson correlation of 4-week post-TAC left ventricular longitudinal strain verses 2-week post-TAC left atrial longitudinal strain. Sample size n=16 for each correlation. Pearson correlation coefficients and corresponding *p*-values displayed. **(D)** ROC analysis using left atrial parameters (max volume: AUC=0.858, *p*<0.0001; ejection fraction: AUC=0.901, p<0.0001; longitudinal strain: AUC=0.908, *p*<0.0001) to predict ventricular ejection fraction above (n=24) or below (n=24) the median (LVEF = 53.7%). **(E)** Principal component analysis scores plot separated by left ventricular ejection fraction above or below the median. **(F)** Ranking of variables in the principal component analysis which contribute most to the separation of groups.

We used principal component analysis (PCA) to highlight variables accounting for the inherent variability in the data. The first two principal components (PCs) explained 64.7% of the variance and visually discriminated between low and high LV EF groups. (Figures 4E-F). Interestingly, LA structural and functional modifications contributed 64.61% to the weight of the PCs (Supplemental Figure 3).

To further evaluate the acute phase of LA remodeling after TAC, we performed a subgroup analysis with imaging at baseline, 3 days post TAC, and 4 weeks post TAC (Figure 5A). Interestingly, LA max volume remained consistent from baseline to 3 days (baseline: 11.2 ± 5.2µL, 3 days: 12.3 ± 3.9µL, *p*=0.856) before significantly increasing to 35.7 ± 16.1µL at 4-weeks post TAC (*p*=0.001; Figure 5B). However, LA ejection fraction (baseline: 53.4 ± 8.2%, 3 days: 23.0 ± 7.5%, *p*<0.001) and strain (baseline: 25.8 ± 9.5%, 3 days: 5.9 ± 2.0%, *p*<0.001) dysfunction were already present three days after induced pressure overload (Figure 5C-D). Furthermore, LV EF was 62.2 ± 5.4% at baseline and remain stable at 56.2 ± 5.1% at 3 days (*p*=0.120) before significantly decreasing to 35.9 ± 3.3% at 4 weeks (Figure 5E). Aortic constriction peak flow velocity was increased in magnitude at 3 days (*p*<0.001) to support a successful TAC (Figure 5F).

**Figure 5:**
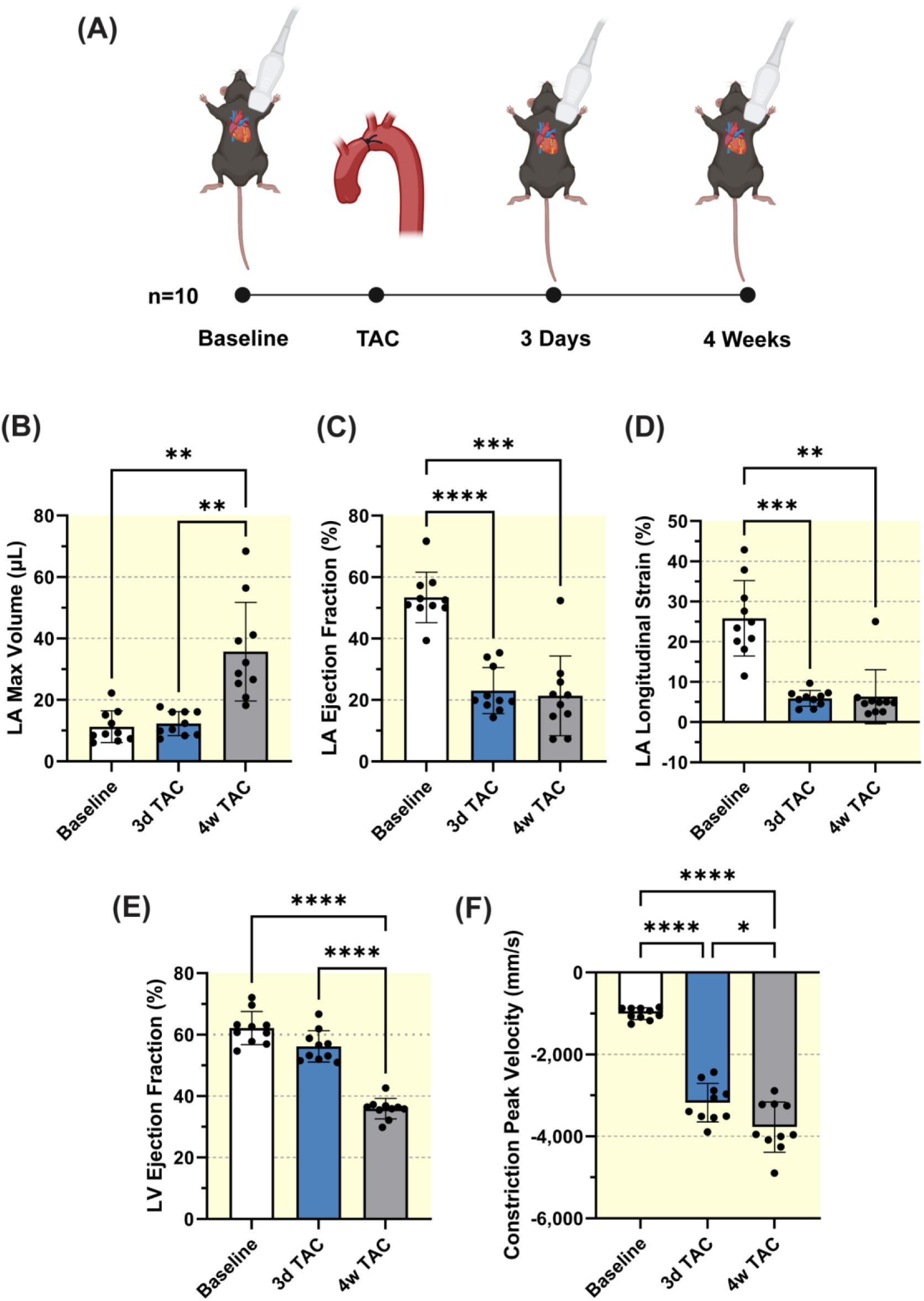
Acute left atrial dysfunction after TAC. **(A)** Study design and timeline. Left atrial **(B)** maximum volume, **(C)** ejection fraction and **(D)** longitudinal strain. **(E)** Left ventricular ejection fraction. **(F)** Constriction peak blood flow velocity. Sample size n=10. Results expressed as mean ± SD. Statistical comparisons conducted with one-way ANOVA with Tukey’s post hoc multiple comparisons tests.

## DISCUSSION

Overall, we report an original approach for LA strain analysis and show that LA dysfunction preceded LV failure. The strong correlation observed between LA strain and subsequent LV loss-of-function suggests that the former is a powerful predictor of heart failure.

Historically, the left atrium has been recognized as a crucial buffer chamber that helps maintain cardiac output and normal LV diastolic pressure (15,16). More recently, measuring LA structure and function has gained emphasis in heart failure management, as LA dysfunction can exacerbate symptoms and is associated with worse outcomes in heart failure patients (6,17). Despite its importance, atrial evaluation is underrepresented in preclinical studies. LA dimensions are a recognized biomarker for studying LV dysfunction across various heart failure models (7,18). However, imaging of the LA in murine models of heart failure remains challenging due to the lack of standardized criteria and consensus, as well as the specific geometry and complex function of the LA ￼￼￼￼￼￼In this study, we demonstrate the feasibility of assessing murine LA dimension using high resolution echocardiography at both early and late stages of pressure overload. We positioned the ultrasound probe in a short-axis view, allowing for clear localization of the LA between the LV base and the pulmonary artery. This orientation enabled us to easily detect both the LA and the left atrial appendage dimensions. Using this technique and in association with histology we confirmed that pressure overload induces structural remodeling in the LA ￼

Anatomically, the LA has a much thinner layer of cardiomyocytes compared to the LV, making it more sensitive to pressure overload and impacts its function. Utilizing specialized speckle-tracking software, we successfully evaluated LA strain trajectories in a murine model of pressure overload. We demonstrated that atrial strain dysfunction and increased LA volume strongly correlate with subsequent LV dysfunction. Furthermore, we hypothesized that LA strain could serve as an early biomarker for heart failure development in this model of transverse aortic constriction. Consistent with previous clinical evidence (11,21), we found that LA dysfunction precedes LV dysfunction. Importantly, using an ROC curve analysis and principal component analysis (PCA), we found that early LA structural and functional modifications can predict the trajectory of LV function following pressure overload. These findings are consistent with recent clinical studies (22,23), which reported that LA dysfunction is highly predictive of cardiovascular outcomes and could be employed as a biomarker to improve risk stratification.

Previous studies have indicated that LA functional impairment can be detected even in the absence of atrial structural changes, suggesting it may be an earlier indicator of cardiac damage, particularly in patients with aortic stenosis (24). Furthermore, some clinical studies have suggested that LA strain dysfunction may precede LA volume adaptations (25,26). Our findings confirm that LA strain dysfunction occurs in the acute phase of pressure overload, preceding abnormal LA geometry, suggesting LA strain is a more sensitive biomarker than volume alone.

The molecular mechanisms underlying structural and functional alterations in the LA are not well understood. In a recent study by Yamaguchi et al., transcriptomic data from a model of pressure overload, two weeks post-intervention, revealed that the most significantly upregulated or downregulated biological processes in the LA involve pathways associated with collagen deposition, hypertrophic and dilated cardiomyopathy, oxidative phosphorylation, and metabolic processes (14). However, the molecular profile of acute LA modifications during the early phase of pressure overload remains unknown. As the LA is highly sensitive to pathologic stimuli such as pressure overload, investigating how gene responses correlate with acute LA strain dysfunction could reveal novel biomarkers and be a crucial area of research in the future. This method offers a promising technique for more accurate and reproducible assessments of LA function in preclinical studies.

### Limitations

LA function was evaluated only in two dimensions, which may have led to under- or overestimation of LA strain. Assessment of LA conduit function was challenging due to the difficulty in distinguishing between the reservoir and contractile phases. Future work could include development of advanced 4D imaging and analysis techniques to accurately evaluate LA function and volume over the cardiac cycle similar to advanced 4D left ventricular analysis techniques (27).

### Conclusion

Our study highlights the potential of murine LA strain analysis as a valuable tool for the early detection of LV dysfunction, offering a promising early biomarker for heart failure development in pressure overload murine models. Future research should focus on the molecular mechanisms underlying these structural and functional alterations in the LA to further refine risk stratification and improve heart failure management.

## METHODS

### Animal Model

Experiments were performed in concordance with European Directive (2010/63/EU) and French laws governing laboratory animal use. The study was approved by the institutional ethics committee (#2023020112386679v3). We used 8-week-old C57BL/6J mice with n=16 in the main cohort and n=5 in the subgroup. The animal habitat was maintained at a constant 21 ± 1°C and 55 ± 1% humidity. Mice had access to food and water at all times.

High resolution ultrasound imaging was performed on a cohort of mice (n=16) at baseline, 2- and 4-weeks post transverse aortic constriction (TAC) (Figure 1A). A subgroup of mice (n=5) and randomly assigned mice from original cohort (n=5) underwent additional evaluation to elucidate the acute effects of TAC on the left atrium. High resolution ultrasound was performed as in the subgroup at baseline, 3-days and 4-weeks post TAC (Figure 5A). Five of the original cohort mice were imaged 3-days post TAC in addition to the baseline, 2-, and 4-week TAC timepoints.

### Transverse Aortic Constriction

Mice were anesthetized with 2.5% isoflurane at 1.0mL/min and a heating pad (Temperature Control Unit HB101/2, Panlab, USA) was used to maintain body temperature at 37°C. Buprenorphine (1 mg/kg) was administered by subcutaneous injection. The animals were secured to a sterile surgical stage in a supine position, and a small incision was made in the neck to visualize the trachea for intubation. The animals were intubated and placed on a ventilator (MiniVent Type 845, Hugo Sachs Elektronik, Germany) with a mixture of room air and 2.5% isoflurane with a respiratory rate of 140 breaths/min and a tidal volume of 225 µL. The ventral thorax hair was removed, and the skin was sterilized with betadine.

As previously described, a closed-chest model was used to induce the transverse aortic constriction (TAC) (28,29). Briefly, a 0.5-1.0 cm transverse incision was made in the left 2^nd^ intercostal space. An electrosurgical unit was used to dissect the pectoralis muscle. A blunt dissection technique was then used to visualize the aortic arch. A small homemade J-hook wire was used to isolate the aortic arch. We placed a 27-gauge needle next to the arch. A 6.0 silk suture was threaded around the needle and aorta between the innominate and left common carotid arteries and a simple suture was made. After confirmation of ligation, the J-hook and excess suture were removed. The intercostal space and the skin were closed using a 5.0 resorbable suture. The isoflurane was stopped, and mice were extubated once spontaneously respiration returned. Mice were allowed to recover under a warming heat pad and were observed further throughout the perioperative period. Following the recovery period, mice were injected twice a day with buprenorphine (1 mg/kg, s.c.).

### Ultrasound Imaging

The Vevo 3100 high-frequency ultrasound system (FUJIFILM VisualSonics) with a 40-MHz center frequency MX550D linear array probe was used to acquire high-resolution 2D and 4D data of the left ventricle and left atrium. We anesthetized the animals as described above without intubation. We removed the ventral thorax hair and secured the mice to the imaging stage in a supine position. The ECG and respiratory rate were monitored throughout the imaging process.

Two-dimensional (2D) echocardiography was performed according to American Physiological Society guidelines for cardiac function measurements in mice (30). Left ventricular volumes and strain were analyzed in the parasternal long-axis view. We acquired four-chamber B-mode images to evaluate ventricular diastolic function using early diastolic strain rate in addition to atrial function. Images of the left atrium and left atrial appendage were obtained in a modified short-axis view (Figure 1B-C).

Four-dimensional (4D) echocardiography was performed at baseline and 4-weeks post-TAC. In the parasternal short axis, a linear step motor with step size 0.13 mm scanning at 400 frames/sec from the apex of the heart to the above the aortic arch was used to acquire 4D data of the left atrium. 4D data not available for all mice.

B-mode and Pulse-Wave Doppler imaging were used to confirm TAC and evaluate constriction blood flow velocity. PW doppler imaging was performed at all time points. We conducted image analysis using Vevo Lab Software 5.9.0 and Vevo Strain 2.0 (FUJIFILM VisualSonics).

### Histology

Baseline (n=4) and 4 week TAC (n=4) hearts were fixed in 4% PFA overnight, dehydrated with a sucrose gradient (final PBS 25% sucrose), frozen in OCT and sectioned (12μm). Sections were permeabilized in PBS 0.1% triton and then blocked for 1h in blocking solution (BS) (PBS 0.1% triton 0.1% BSA). Sections were then incubated overnight with Goat Anti-Type I Collagen (Southern Biotech) 1/200 in BS, washed in PBS and incubated for 2h with secondary antibody and DAPI in BS, then washed and mounted with Fluoromount-G (Thermo Fisher Scientific). Sections were imaged using a slide scanner (Zeiss Axioscan) and Type I Collagen staining was quantified in atria using ImageJ software. At least 2 equivalent sections of left atria were analyzed per mouse and per condition.

### Statistical Analysis

All statistical analysis was performed using Prism 10.2.3 (GraphPad Software). The threshold for statistical significance was set at *p* < 0.05. Baseline, 3 days, 2- and 4-weeks post-TAC parameters are shown as mean ± standard deviation (SD). A one-way ANOVA with Tukey post-hoc multiple pairwise comparison’s test was used to compare parameters across timepoints. Correlation between parameters was described using Pearson correlation coefficients if data was normally distributed as indicated by a Shapiro – Wilk test; otherwise, a Spearman’s correlation coefficient and associated *p*-value were calculated.

We performed a receiver operating characteristic (ROC) analysis to evaluate discriminative performance of left atrial function parameters on left ventricular function. The ROC groups were determined based on the median LV ejection fraction. LA max volume, LA ejection fraction, and LA strain were then used to discriminate between LV ejection fraction values above or below the median.

A principal component analysis (PCA) was performed to elucidate meaningful variables. The analysis included LA and LV function parameters from baseline, 2-, and 4-weeks post TAC. A standardized PCA method was used to account for variable scale differences. Principal components (PCs) were selected based on a parallel analysis with 1,000 simulations and a 95% percentile level. The first two PCs are plotted to visualize how PCA discriminates between groups of LV ejection fraction above or below the median given all the data. The contribution of the variables to PC1 is shown to evaluate variable importance. All material submitted conforms to good publishing practice in physiology (31).

## Ethics Approval

The study was approved by the institutional ethics committee (#2023020112386679v3).

## Data Availability Statement

The data that support the findings of this study are available from the corresponding author upon reasonable request.

## Disclaimers

All authors have read the journal’s policy on disclosure of potential conflicts of interest. C. J. Goergen serves on the Scientific Advisory Board for FUJIFILM VisualSonics Inc. None of the other authors have conflicts of interest, financial or otherwise, to disclose. Disclaimers: FUJIFILM VisualSonics Inc. had no role in the study’s design, execution, interpretation, or writing.

## Author Contributions

JPS designed, planned, and performed the strain analysis and most of the experimental work including data analysis; prepared the figures; and drafted the manuscript. TMM, CJG, and PS initiated, designed, and provided continuous guidance in the conceptual design of the work, data interpretation, reviewed and edited the manuscript. All the authors edited and approved the manuscript.

## Acknowledgements

We gratefully thank the staff for animal housing (PhyMedExp). We acknowledge the technical support provided by Salomé Bélec from Imagerie du Petit Animal de Montpellier (IPAM) for accessing high-resolution ultrasound (LRQA Iso9001; France Life Imaging (grant ANR-11-INBS-0006); IBISA; Leducq Foundation (RETP), I-Site Muse).

---

**Supplemental Figure 1:** Estimated two-dimensional atrial volumes correlate with measured four-dimensional volumes. **(A)** Representative left atrial volume at baseline and 4 weeks post-TAC. Scale bar represents 3mm **(B)** Left atrial 4D volumes over time. Sample size n=16 for baseline and n=11 for 4 weeks post-TAC. Results expressed as mean ± SD. Statistical comparison conducted with non-paired studentized t-test. **(C)** Spearman nonparametric correlation between 4D and 2D left atrial volumes. Sample size n=27. Correlation coefficient and corresponding *p*-value displayed.

**Supplemental Figure 2:** Correlation of left atrial and left ventricular function parameters. **(A)** Correlation between left atrial and left ventricular function parameters from all time points, n=48. **(B)** Correlation between left atrial function parameters at 2 weeks post-TAC and left ventricular function parameters at 4 weeks post-TAC, n=16. Pearson correlation coefficients, or Spearman coefficients if data non-normally distributed as determined by Shapiro-Wilk test, displayed. Colors depict levels of statistical significance from corresponding *p*-values.

**Supplemental Figure 3:** Proportion of variance in LV ejection fraction accounted for by each successive principal component.

