## Supplemental figure for "Left Atrial Strain as a Predictor of Cardiac Dysfunction in a Murine Model of Pressure Overload"

### Supplemental Figure 1:

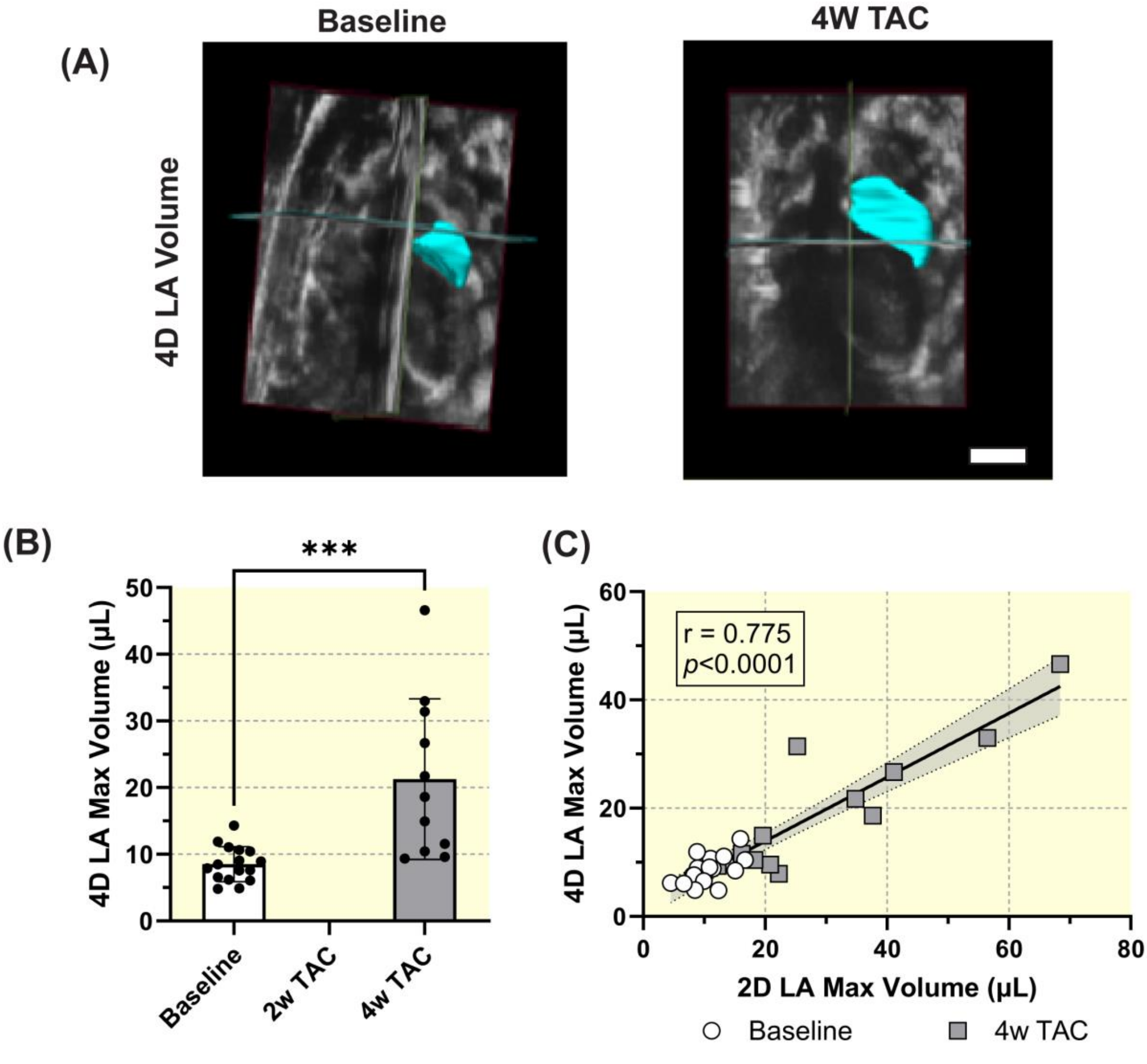

### Supplemental Figure 2:

**(A)**

| All Time Points Correlation | LA Max Volume | LA Min Volume | LA Area | LA Stroke Volume | LA Ejection Fraction | LA Longitudinal Strain | LA Circumferential Strain | LA Fractional Area Change | Peak Flow Velocity | LV End-Diastoli Volume | LV Peak-Systolic Volume | LV Stroke Volume | LV Ejection Fraction | LV Longitudinal Strain | LV Early Diastolic Strain Rate | LV Circumferential Strain |
|---|---|---|---|---|---|---|---|---|---|---|---|---|---|---|---|---|
| LA Max Volume | 1.000 | 0.956 | 0.889 | 0.414 | -0.699 | -0.594 | -0.756 | -0.603 | -0.575 | 0.430 | 0.609 | -0.253 | -0.690 | 0.594 | -0.276 | 0.350 |
| LA Min Volume | 0.956 | 1.000 | 0.877 | 0.220 | -0.855 | -0.706 | -0.842 | -0.765 | -0.622 | 0.415 | 0.605 | -0.287 | -0.721 | 0.659 | -0.327 | 0.346 |
| LA Area | 0.889 | 0.877 | 1.000 | 0.317 | -0.681 | -0.667 | -0.673 | -0.597 | -0.708 | 0.493 | 0.670 | -0.180 | -0.715 | 0.750 | -0.432 | 0.253 |
| LA Stroke Volume | 0.414 | 0.220 | 0.317 | 1.000 | 0.212 | 0.258 | 0.027 | 0.282 | -0.014 | 0.111 | 0.152 | -0.060 | -0.153 | 0.007 | 0.000 | -0.134 |
| LA Ejection Fraction | -0.699 | -0.855 | -0.681 | 0.212 | 1.000 | 0.802 | 0.859 | 0.939 | 0.576 | -0.378 | -0.583 | 0.284 | 0.599 | -0.681 | 0.366 | -0.453 |
| LA Longitudinal Strain | -0.594 | -0.706 | -0.667 | 0.258 | 0.802 | 1.000 | 0.648 | 0.755 | 0.622 | -0.292 | -0.525 | 0.224 | 0.563 | -0.643 | 0.387 | -0.437 |
| LA Circumferential Strain | -0.756 | -0.842 | -0.673 | 0.027 | 0.859 | 0.648 | 1.000 | 0.688 | 0.521 | -0.451 | -0.600 | 0.197 | 0.607 | -0.636 | 0.348 | -0.346 |
| LA Fractional Area Change | -0.603 | -0.765 | -0.597 | 0.282 | 0.939 | 0.755 | 0.688 | 1.000 | 0.558 | -0.307 | -0.516 | 0.235 | 0.551 | -0.588 | 0.323 | -0.402 |
| Peak Flow Velocity | -0.575 | -0.622 | -0.708 | -0.014 | 0.576 | 0.622 | 0.521 | 0.558 | 1.000 | -0.336 | -0.614 | 0.289 | 0.744 | -0.759 | 0.594 | -0.207 |
| LV End-Diastoli Volume | 0.430 | 0.415 | 0.493 | 0.111 | -0.378 | -0.292 | -0.451 | -0.307 | -0.336 | 1.000 | 0.835 | 0.428 | -0.507 | 0.511 | -0.261 | 0.231 |
| LV Peak-Systolic Volume | 0.609 | 0.605 | 0.670 | 0.152 | -0.583 | -0.525 | -0.600 | -0.516 | -0.614 | 0.835 | 1.000 | -0.012 | -0.885 | 0.748 | -0.428 | 0.424 |
| LV Stroke Volume | -0.253 | -0.287 | -0.180 | -0.060 | 0.284 | 0.224 | 0.197 | 0.235 | 0.289 | 0.428 | -0.012 | 1.000 | 0.457 | -0.297 | 0.270 | -0.227 |
| LV Ejection Fraction | -0.690 | -0.721 | -0.715 | -0.153 | 0.599 | 0.563 | 0.607 | 0.551 | 0.744 | -0.507 | -0.885 | 0.457 | 1.000 | -0.760 | 0.484 | -0.424 |
| LV Longitudinal Strain | 0.594 | 0.659 | 0.750 | 0.007 | -0.681 | -0.643 | -0.636 | -0.588 | -0.759 | 0.511 | 0.748 | -0.297 | -0.760 | 1.000 | -0.694 | 0.347 |
| LV Early Diastolic Strain Rate | -0.276 | -0.327 | -0.432 | 0.000 | 0.366 | 0.387 | 0.348 | 0.323 | 0.594 | -0.261 | -0.428 | 0.270 | 0.484 | -0.694 | 1.000 | -0.115 |
| LV Circumferential Strain | 0.350 | 0.346 | 0.253 | -0.134 | -0.453 | -0.437 | -0.346 | -0.402 | -0.207 | 0.231 | 0.424 | -0.227 | -0.424 | 0.347 | -0.115 | 1.000 |

| | | |
|---|---|---|
| $p<0.05$ | $p<0.01$ | $p<0.001$ |

**(B)**

| 2w LA vs 4w LV Correlation | LA Max Volume | LA Min Volume | LA Area | LA Stroke Volume | LA Ejection Fraction | LA Longitudinal Strain | LA Circumferential Strain | LA Fractional Area Change | Peak Flow Velocity | LV End-Diastoli Volume | LV Peak-Systolic Volume | LV Stroke Volume | LV Ejection Fraction | LV Longitudinal Strain | LV Early Diastolic Strain Rate | LV Circumferential Strain |
|---|---|---|---|---|---|---|---|---|---|---|---|---|---|---|---|---|
| LA Max Volume | 1.000 | 0.985 | 0.897 | 0.229 | -0.706 | -0.773 | -0.638 | -0.676 | -0.279 | 0.278 | 0.341 | -0.224 | -0.306 | 0.629 | -0.426 | 0.612 |
| LA Min Volume | 0.985 | 1.000 | 0.876 | 0.174 | -0.776 | -0.801 | -0.694 | -0.685 | -0.300 | 0.193 | 0.244 | -0.235 | -0.250 | 0.591 | -0.447 | 0.582 |
| LA Area | 0.897 | 0.876 | 1.000 | 0.250 | -0.609 | -0.773 | -0.538 | -0.541 | -0.315 | 0.414 | 0.438 | -0.218 | -0.371 | 0.632 | -0.279 | 0.679 |
| LA Stroke Volume | 0.229 | 0.174 | 0.250 | 1.000 | 0.232 | 0.185 | 0.218 | 0.207 | 0.344 | -0.194 | -0.264 | 0.170 | 0.326 | -0.167 | 0.227 | -0.319 |
| LA Ejection Fraction | -0.706 | -0.776 | -0.609 | 0.232 | 1.000 | 0.930 | 0.929 | 0.869 | 0.354 | -0.162 | -0.278 | 0.266 | 0.315 | -0.547 | 0.504 | -0.629 |
| LA Longitudinal Strain | -0.773 | -0.801 | -0.773 | 0.185 | 0.930 | 1.000 | 0.889 | 0.748 | 0.564 | -0.183 | -0.315 | 0.302 | 0.389 | -0.553 | 0.450 | -0.642 |
| LA Circumferential Strain | -0.638 | -0.694 | -0.538 | 0.218 | 0.929 | 0.889 | 1.000 | 0.658 | 0.355 | -0.166 | -0.283 | 0.270 | 0.333 | -0.534 | 0.407 | -0.583 |
| LA Fractional Area Change | -0.676 | -0.685 | -0.541 | 0.207 | 0.869 | 0.748 | 0.658 | 1.000 | 0.178 | -0.153 | -0.180 | 0.075 | 0.146 | -0.337 | 0.361 | -0.483 |
| Peak Flow Velocity | -0.279 | -0.300 | -0.315 | 0.344 | 0.354 | 0.564 | 0.355 | 0.178 | 1.000 | -0.210 | -0.419 | 0.471 | 0.608 | -0.451 | 0.424 | -0.495 |
| LV End-Diastoli Volume | 0.278 | 0.193 | 0.414 | -0.194 | -0.162 | -0.183 | -0.166 | -0.153 | -0.210 | 1.000 | 0.888 | 0.137 | -0.578 | 0.585 | -0.027 | 0.655 |
| LV Peak-Systolic Volume | 0.341 | 0.244 | 0.438 | -0.264 | -0.278 | -0.315 | -0.283 | -0.180 | -0.419 | 0.888 | 1.000 | -0.335 | -0.884 | 0.785 | -0.284 | 0.829 |
| LV Stroke Volume | -0.224 | -0.235 | -0.218 | 0.170 | 0.266 | 0.302 | 0.270 | 0.075 | 0.471 | 0.137 | -0.335 | 1.000 | 0.719 | -0.492 | 0.557 | -0.443 |
| LV Ejection Fraction | -0.306 | -0.250 | -0.371 | 0.326 | 0.315 | 0.389 | 0.333 | 0.146 | 0.608 | -0.578 | -0.884 | 0.719 | 1.000 | -0.772 | 0.451 | -0.800 |
| LV Longitudinal Strain | 0.629 | 0.591 | 0.632 | -0.167 | -0.547 | -0.553 | -0.534 | -0.337 | -0.451 | 0.585 | 0.785 | -0.492 | -0.772 | 1.000 | -0.584 | 0.877 |
| LV Early Diastolic Strain Rate | -0.426 | -0.447 | -0.279 | 0.227 | 0.504 | 0.450 | 0.407 | 0.361 | 0.424 | -0.027 | -0.284 | 0.557 | 0.451 | -0.584 | 1.000 | -0.412 |
| LV Circumferential Strain | 0.612 | 0.582 | 0.679 | -0.319 | -0.629 | -0.642 | -0.583 | -0.483 | -0.495 | 0.655 | 0.829 | -0.443 | -0.800 | 0.877 | -0.412 | 1.000 |

| | | |
|---|---|---|
| $p<0.05$ | $p<0.01$ | $p<0.001$ |

### Supplemental Figure 3:

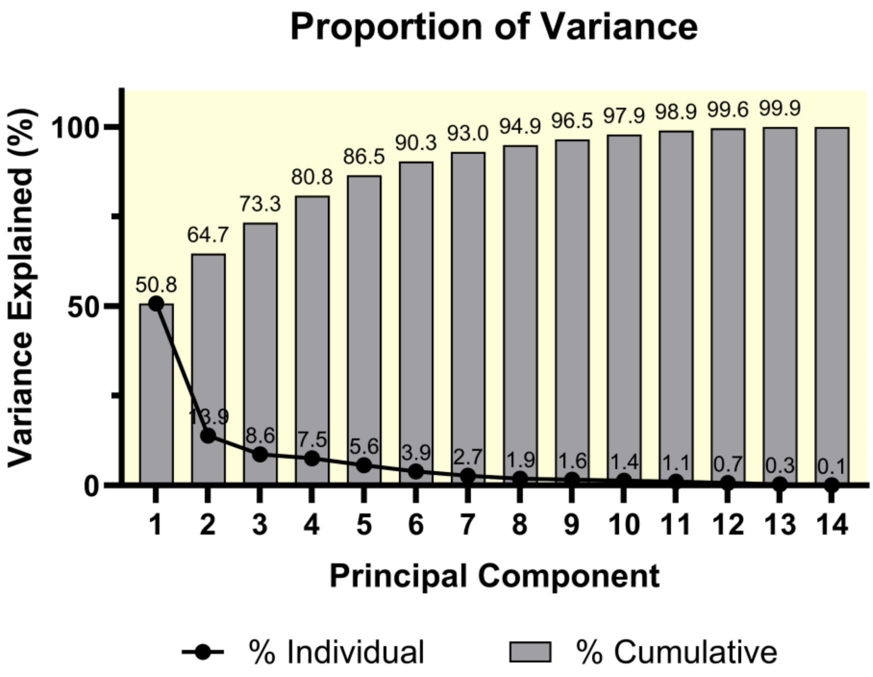
